# Differential interactions determine anisotropies at interfaces of RNA-based biomolecular condensates

**DOI:** 10.1101/2024.08.19.608662

**Authors:** Nadia A. Erkamp, Mina Farag, Yuanxin Qiu, Daoyuan Qian, Tomas Sneideris, Tingting Wu, Timothy J. Welsh, Hannes Ausserwӧger, Tommy J. Krug, David A. Weitz, Matthew Lew, Tuomas P. J. Knowles, Rohit V. Pappu

## Abstract

Biomolecular condensates form via macromolecular phase separation, giving rise to coexisting phases delineated by interfaces. Here, we characterize the structures of interfaces formed by phase separation driven by heterotypic interactions in ternary mixtures of two types of RNA molecules and polyethylene glycol. We find that purine-rich RNAs are scaffolds that drive phase separation via strong heterotypic interactions. Conversely, pyrimidine-rich RNA molecules are defined by weaker heterotypic interactions. They function as adsorbents that accumulate at and wet the interfaces of coexisting phases formed by phase separation of scaffolds. Our computations predict that scaffolds and adsorbents have different non-random orientational preferences at interfaces. We tested these predictions using single-molecule super resolution imaging that tracks the motions of fluorogenic probes that are bound to RNA molecules. Motions parallel to the interface were found to be faster than motions perpendicular to the interface. These findings support previous predictions regarding anisotropies of motions at interfaces.

---

Biomolecular condensates are compositionally distinct membraneless bodies that form via phase separation of macromolecules from bulk solutions ^1,2^. Condensates are two- or multi-phase systems that provide spatial and temporal control and organization over biochemical reactions involving proteins and nucleic acids ^1,2^. There is growing interest in developing synthetic condensates with bespoke cellular functions ^3–5^. This is being driven by numerous efforts that have advanced our understanding of the physico-chemical principles that drive the formation and dissolution of condensates in live cells ^6,7^ and *in vitro* ^8^. In live cells, condensates comprise several distinct types of macromolecules ^9^. Further, within dense phases of condensates, one often observes a hierarchical organization of the constituent macromolecules, and this has been reported for stress granules ^10–12^, nucleoli ^13,14^, nuclear speckles ^15^, and the mitochondrial nucleoid ^16^. Spatially organized condensates were also shown to arise in simple mixtures ^17^ including ternary systems comprising two different RNA molecules and an arginine-rich peptide ^18^.

Increased attention is being paid to interfaces of condensates ^19,20^. A recent computational study predicted that in single component condensates, the polymers that drive condensation, referred to as scaffolds, prefer non-random, orthogonal orientations with respect to the plane of the interface ^19^. The solidification of condensates can be nucleated from the interface ^21–24^. Additionally, structured interfaces can affect condensate fusion ^25^, catalyze redox reactions ^26^ and generate resistance for molecules moving from dilute to dense phase or vice versa ^27^. Notably, in cells, long or small non-coding RNA molecules feature prominently in several nuclear bodies such as nuclear speckles ^15^, peri-nucleolar compartments ^28^, and paraspeckles ^29^. RNA molecules can be scaffolds that drive phase separation from the bulk or they can adsorb onto interfaces of condensates formed by scaffolds. Understanding why specific RNA molecules adsorb is relevant for understanding how these condensates function in cells ^20^, and for designing *de novo* condensates based on protein-RNA mixtures ^5,20^.

Here, we investigate the principles underlying spatial organization in condensates formed by ternary mixtures using a model system that comprises a biocompatible polymer, namely polyethylene glycol (PEG)^30^, and pairs of homopolymeric, non-base-pairing RNA molecules such as poly-adenine (poly-rA), which is a poly-purine, and poly-cytosine (poly-rC), which is a poly-pyrimidine. Our investigations are motivated by a general framework anchored in the Gibbs adsorption isotherm ^31^, and the physics of adsorption-mediated wetting transitions ^32^. We study adsorption in condensates formed by ternary mixtures and investigate the effects of polymer interaction strengths, concentrations, lengths, and stiffness on adsorption. Then, we quantify the orientational preferences of molecules at the interface that leads to the distinction of condensate components being scaffolds versus adsorbents. We report results from single-molecule super resolution imaging that were used to study the movement of molecules near the interface. With respect to the interface, we observe significantly faster motion in the parallel direction than in the perpendicular direction. Finally, we discuss the broader implications of our findings for the design of *de novo* condensates with specific spatial organization and interfacial properties.

## Results

### Pyrimidine-rich RNA undergoes adsorption and wetting transitions

Heterotypic PEG-RNA interactions provide the driving forces for forming condensates in the model ternary systems ^30,33^. Increasing the concentration of PEG alters the chemical potential of PEG and promotes phase separation. Likewise, increasing the concentration of NaCl weakens electrostatic repulsions between RNA molecules, and this also promotes phase separation. We first obtained a coarse mapping of the overall phase behaviors as a function of NaCl and PEG concentrations. Poly-rA undergoes phase separation in the presence of sufficient PEG and NaCl (**Fig. 1a-b**). The amount of NaCl required to drive phase separation decreases as the concentration of PEG increases, and the slope of the phase boundary becomes zero above 3 w/w% of PEG. For these concentrations of PEG, the poly-rA + PEG mixture undergoes phase separation for all NaCl concentrations that are above ∼0.5 M.

**Figure 1:**
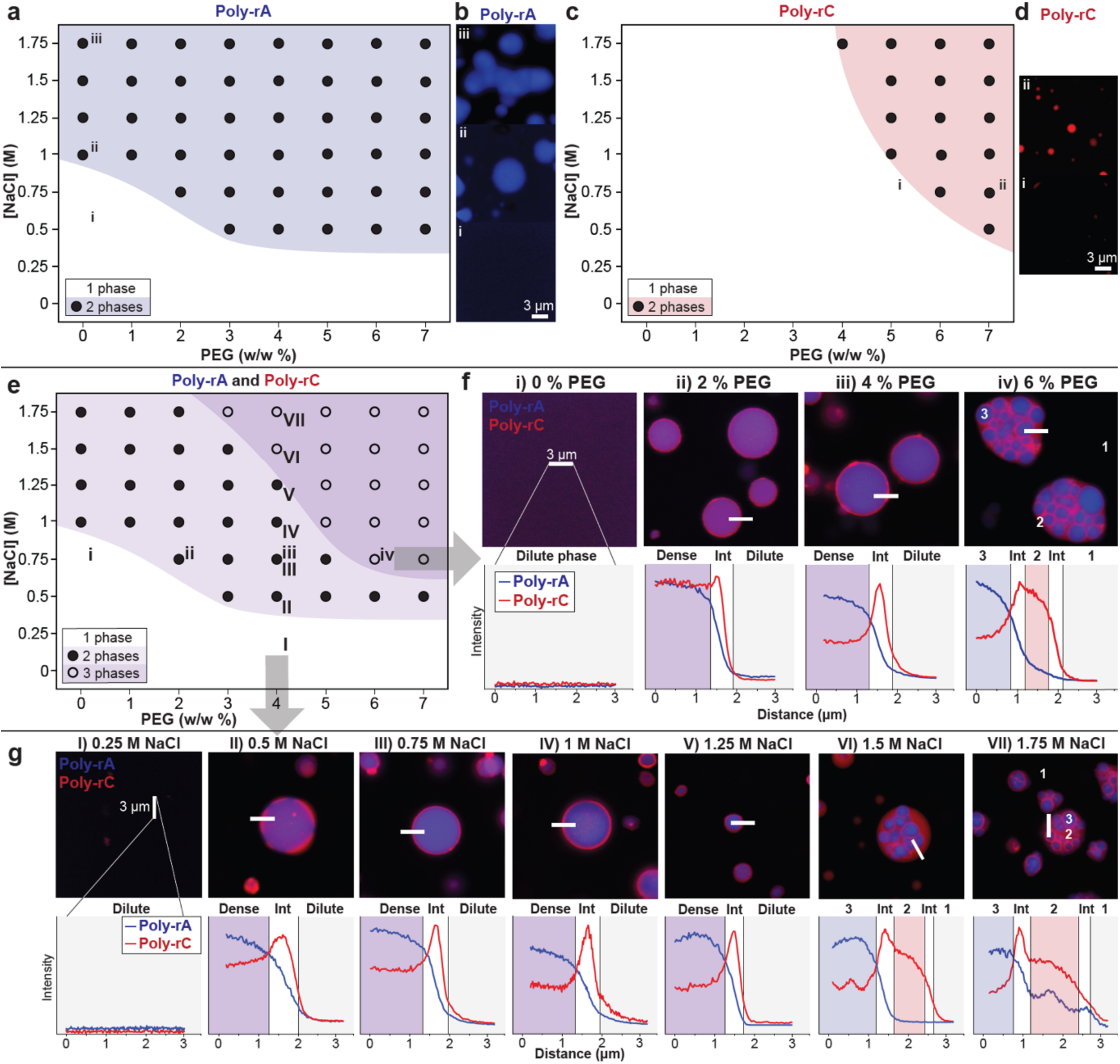
Phase separation and adsorption of Poly-rA and -rC. **a** Poly-rA (1 g/L) undergoes phase separation in the presence of high concentrations of NaCl and PEG, **b** forming spherical (ii) or irregularly-shaped condensates. **c** poly-rC (1 g/L) phase separation requires higher concentrations of NaCl and PEG **d** to form spherical condensates. **e** The dense phases in a mixture of poly-rA (1 g/L) and poly-rC (1 g/L) show a range condensate architectures. At low concentrations of PEG (i of **f**) or low concentrations of NaCl (I of **g**) the dense phase is homogenous Increasing either the concentration of PEG or NaCl, poly-rA forms condensates and some poly-rC is recruited into it (ii of **f**, II or **g**). Notably, poly-rC adsorbs to the interface. The extent of adsorption increases with increasing concentrations of PEG (iii of **f**) or NaCl (III, IV, V of **g**). At higher concentrations of PEG, for fixed NaCl, or higher concentrations of NaCl for fixed PEG, the adsorbed poly-rC undergoes phase separation leading to a poly-rA-rich phase coexisting with a poly-rC-rich phase. Adsorption of poly-rC around the poly-rA dense phase is still observed (iv of **f**, VI and VII of **g**). All scale bars in the confocal images represent 3 μm.

While poly-rC also undergoes phase separation, higher amounts of PEG and NaCl are required (**Fig. 1c-d**). This shows that poly-rA drives phase separation through stronger interactions when compared to poly-rC^18,34^. We reason that the double rings of poly-purines make them more hydrophobic and allow for stronger π-stacking than poly-pyrimidines ^35–37^ in these systems where base-pairing is not possible. Stronger interactions in purine-rich systems can also engender dynamically arrested phase separation ^16,34^, giving rise to irregular mesoscale structures that are dominated by long-lived intermolecular crosslinks. The resultant percolated networks were observed for poly-rA and not for poly-rC (**Fig. 1b, d**) – an observation that is consistent with recent reports ^37^.

Next, we mapped phase boundaries for ternary mixtures of PEG plus a 1:1 ratio of poly-rA and poly-rC (**Fig. 1e-g**). At 0% PEG and 0.75 M NaCl, we do not observe phase separation (**i, Fig 1f**). Increasing the PEG concentration to 2% w/w, leads to crossing of the phase boundary and formation of a poly-rA-rich phase that coexists with a dilute phase (**ii, Fig 1f**). The poly-rC molecules partition into and adsorb onto the interface of the poly-rA-rich phase. The extent of adsorption increases with increasing concentration of PEG (**iii, Fig 1f**). When the concentration threshold for forming poly-rC-rich phases is crossed, we observe the formation of three coexisting phases: a dilute phase that coexists with poly-rA- and poly-rC-rich phases, with an interface between the latter two phases (**iv, Fig 1f**). Notably, poly-rC still adsorbs to the poly-rA interfaces. The transition from two coexisting phases with adsorption to three coexisting phases is known as a wetting transition ^32^.

We observed similar wetting transitions when varying the concentration of NaCl. In 4% PEG and 0.25 M NaCl, the system forms a homogeneous solution (**I, Fig 1g**). As the NaCl concentration increases. In the presence of 0.5 M NaCl, poly-rA forms condensates and poly-rC partitions into and adsorbs onto the interface of the poly-rA-rich phase (**II, Fig 1g**). The term adsorption refers to the density of poly-rC at the interface being higher than its density inside the poly-rA-rich phase. Between 0.75 M and 1.25 M NaCl, we observe what Kaplan et al.^32^, refer to as complete wetting whereby poly-rC forms a well-defined, micron-scale layer at the interface (**III-V, Fig 1g**). At 1.5 M NaCl and above, we observe a distinct poly-rC-rich phase forms that coexists with a poly-rA-rich phase (**VI VII, Fig 1g**). the cascade of transitions we observe for poly-rC can be explained as follows: At low salt concentrations, poly-rC is an adsorbent, forming favorable interactions with PEG molecules at the interface. Increasing the salt concentration beyond the levels that saturate the interface, leads to the formation of two coexisting phases defined by favorable poly-rA and PEG interactions (the poly-rA-rich phase), and less favorable poly-rC-PEG interactions, which gives rise to the poly-rC-rich phase. The line scans show the coexistence of poly-rA-rich and poly-rC-rich phases separated by a poly-rC-rich interface (**VI VII, Fig 1g**). Note that increasing the salt concentration enhances the heterotypic PEG-poly-rC interactions and the homotypic interactions between poly-rC.

To assess the transferability of our findings to other systems, we have explored a range of RNA homopolymers and their resulting phase behaviors. Specifically, we explored the ability of rA_20_, rA_80_, poly-rA, rC_20_, rC_80_, poly-rC, rU_20_, rU_80_, poly-rU and rG_20_ to adsorb onto condensates of poly-rA, poly-rC and/or poly-rU with PEG (**Ext. Data Fig. 1**, 2). RNA molecules with long guanine tracts can fold into G-quadruplexes. These structures stack on top of one another to form aggregates ^38^. For this reason, rG_80_ and poly-rG were excluded from this investigation. Out of the 35 unique combinations that were explored, we observed a clear trend whereby poly-pyrimidines (rC, rU) adsorb onto dense phases rich in poly-purines. This observation is consistent with the fact that the poly-rA, which is a poly-purine, is a stronger driver of phase separation with PEG (**Fig. 1a**) when compared poly-rC (**Fig. 1c**). The interaction strengths between the RNA and PEG are influenced by whether the RNA is a poly-pyrimidine or poly-purine, given the difference in hydrophobicity and ability to undergo π-stacking ^35^. Thus, our results suggest that in ternary mixtures, the differences in the relative interaction strengths determine which RNAs become the scaffolds and which RNAs are adsorbents that undergo concentration-dependent wetting transitions.

**Figure 2:**
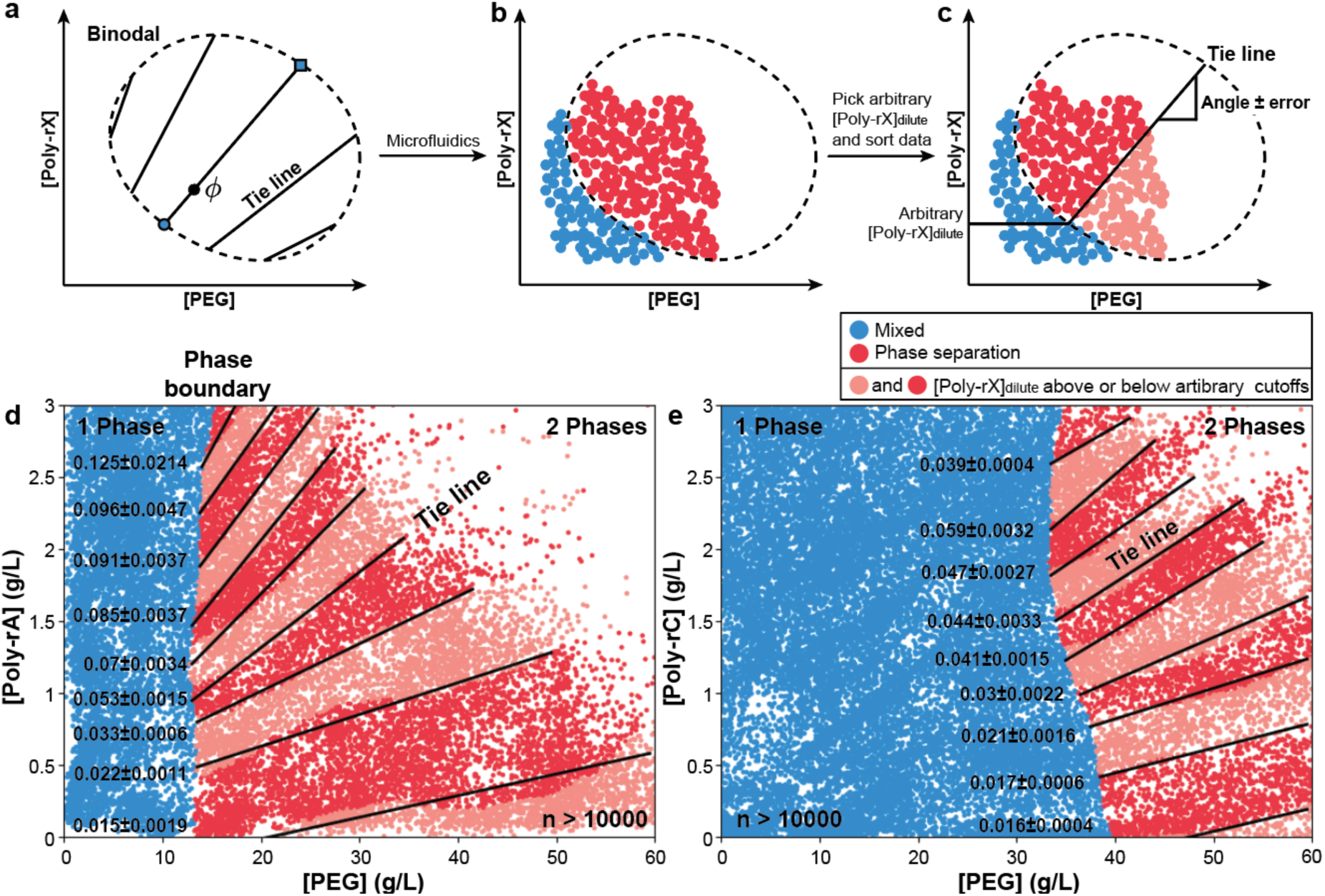
Measurements of phase boundaries and tie line proxies using microfluidics. **a**Tie lines show how a sample with total composition φ separates into a dilute phase (blue circle) and dense phase (blue square). Tie line angles quantify the repulsion versus attraction of molecules in the two phases (**Ext. Data Fig. 3**). **b** Using microfluidics (**Ext. Data Fig. 4**), we can find a phase boundary (dashed line) and obtain proxies for tie lines using dilute phase concentrations. Phase diagrams of **d** poly-rA (1 g/L) and **e** poly-rC (1 g/L) generated using a microfluidic platform. Solid lines show the tie line proxies. Phase separation requires less PEG for poly-rA (**d**), than for poly-rC (**e**). Full lines show a tie line proxy. The tie line proxy of poly-rA with PEG (**c**) is steeper than that of poly-rC with PEG (**d**). This indicates that PEG interacts stronger with poly-rA than with poly-rC.

### Adsorbents interact weakly with scaffolds

Next, we compared the relative interaction strengths between PEG and different RNAs by comparing phase diagrams of different ternary mixtures. Specifically, we compare both the location of the phase boundary (binodal, dashed line) and tie lines (solid lines) (**Fig. 2a**). A sample with a total composition of φ_total_ separates into a dilute phase (blue circle with composition φ_dilute_) and dense phase (blue square with composition φ_dense_). The tie line is the line connecting the points φ_dilute_, φ_total_, and φ_dense_. The angle subtended at the intersection of the tie line with the low concentration arm of the phase boundary helps quantify the relative inclusion or exclusion of molecules in the dilute and dense phases ^33^. The slopes of tie lines quantify the relative contributions of homotypic versus heterotypic interactions in the RNA and PEG system. A positive slope indicates that favorable heterotypic interactions drive the formation of condensates, whereas a negative slope indicates that heterotypic interactions are unfavorable. We mapped phase diagrams using a previously established microfluidics platform ^33,39^, creating ≈10^4^ samples with varying amounts of RNA and PEG. The approach known as PhaseScan scans and sorts of a wide cross section of samples, annotating these as homogeneous (blue) or two-phase systems containing condensates (red). The totality of measurements across the entire spectrum of PEG and RNA concentrations that were scanned helps in mapping the boundary between the one-phase and two-phase regimes (**Fig. 2b**). Additionally, we use the concentrations of RNA in the coexisting dilute phase to infer proxies for tie lines (**Fig. 2c**). This is a way to leverage data from high throughput scanning of phase behaviors to estimate a useful proxy for the actual tie line (**Ext. Data Fig. 3**).

**Figure 3:**
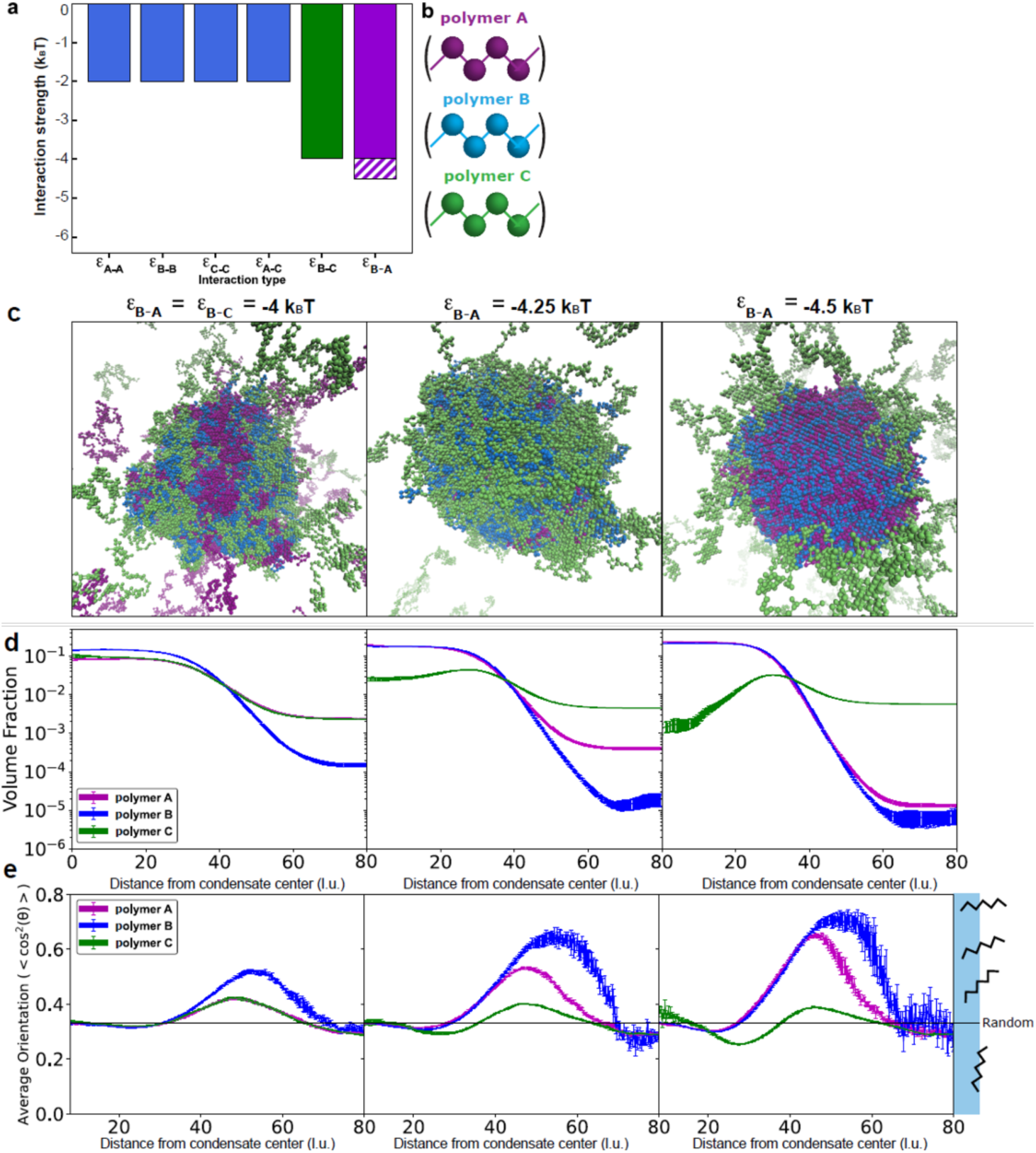
Pairwise interaction strengths and relative concentrations determine condensate architectures and interfacial features. **a**The interaction strengths between polymer beads used in the simulations. The interaction between beads of polymers A and B was varied from –4 to –4.5. **b** Molecules A, B, C represent poly-rA, PEG and poly-rC respectively. **c** Representative snapshots from LaSSI simulations showing distinct condensate architectures obtained by varying the interaction strength between beads of polymers A and B, |ε_B-A_| = –4, –4.25, –4.5. For this same titration, we show **d** the radial density profiles and **d** the average cos^2^8 values, which indicate the average orientation of molecules with respect to the center of the condensate (Supplementary methods). If a given datapoint had an error greater than 0.1, then that datapoint was omitted. In **d** and **e**, error bars indicate the standard errors about the mean across three replicates.

Analysis of our microfluidics data (**Ext. Data Fig. 4**) helps us delineate low concentration arms of phase boundaries for the poly-rA plus PEG (**Fig. 2d**) and poly-rC plus PEG systems (**Fig. 2e**). As expected, poly-rA undergoes phase separation at lower PEG concentrations than poly-rC. The slopes of the tie line proxies are positive for both mixtures, indicating favorable heterotypic interactions between RNA and PEG^30,33^ and partitioning of both PEG and RNA into the dense phases. Notably, tie line proxies for poly-rA with PEG are steeper than for poly-rC with PEG. This implies that the favorable interactions between poly-rA and PEG are stronger than the favorable interactions for poly-rC with PEG.

**Figure 4:**
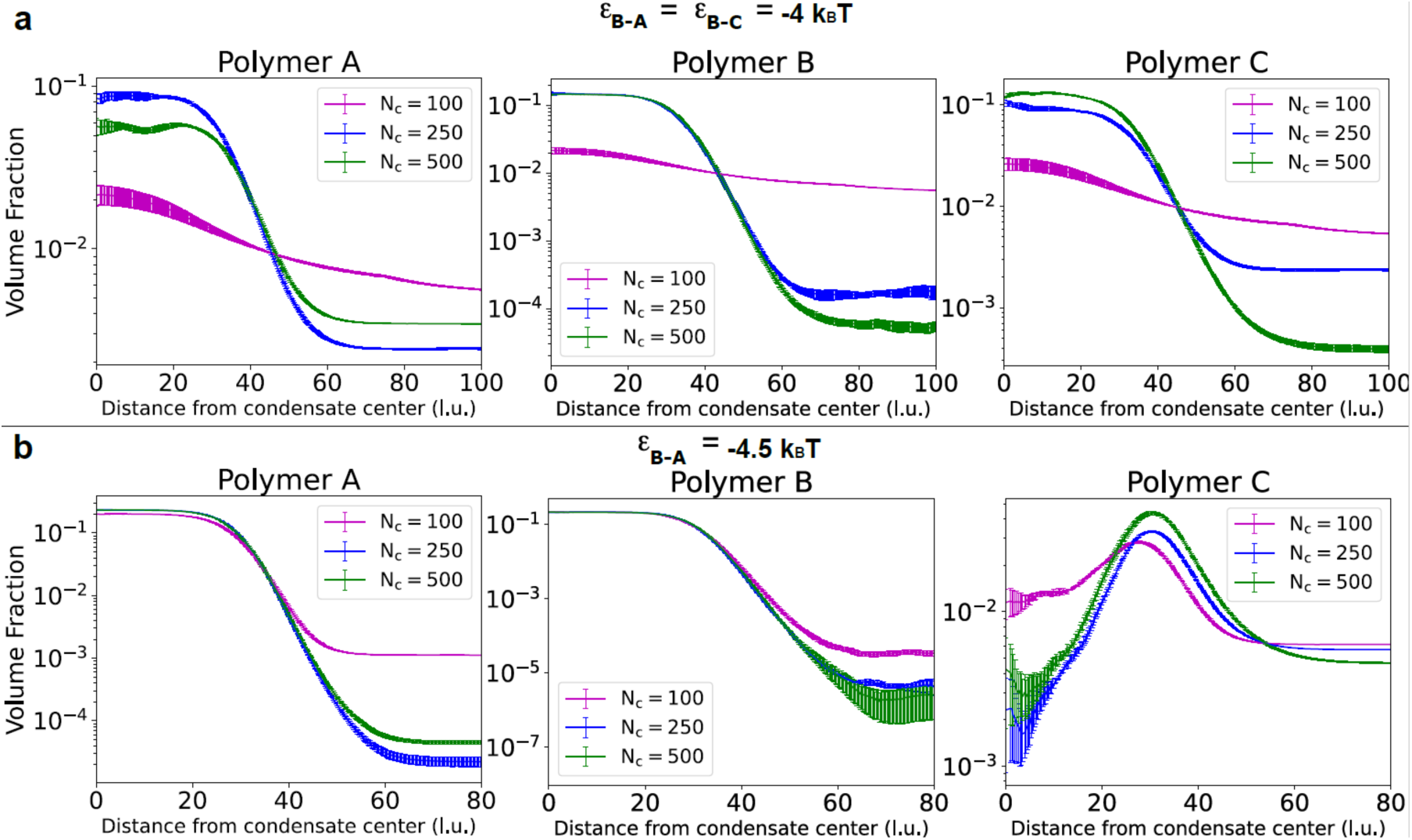
Polymer length has a limited effect on condensate architecture in comparison to pairwise interaction strengths. The interaction between beads of polymers A and B was set to **a** |ε_B-A_| = –4, or **b** |ε_B-A_| = – 4.5. The volume fractions for the three polymers are shown as a function of the distance from the center of the condensate (like Fig. 3d). We observe that molecule length affects partitioning into dilute or dense phase. Notably, varying length does not cause polymer C to **a** adsorb when it previously did not, or **b** stop adsorbing when it previously did. The interaction strength between the monomers has a much more extensive effect on condensate architecture and interface organization than the length of the polymer.

Taken together with the data in **Fig. 1**, we arrive at a phenomenological explanation of the adsorption data. Poly-rA and PEG interact favorably, and this drives phase separation with the dense phases being enriched in both species even at moderate PEG concentrations. The favorable interactions between poly-rC and PEG are weaker when compared with poly-rA. This allows poly-rC to partition into the poly-rA + PEG rich phase. Partitioning of poly-rC into poly-rA-rich phases creates a competition between poly-rA and poly-rC. The latter cannot compete effectively for interactions within the dense phase, and as a result, the partitioning is weak, and it adsorbs onto the interface of the poly-rA-rich phase. At the interface, poly-rC can interact with PEG that is not engaged in interactions with poly-rA.

Overall, the relative strengths of the RNA-PEG interactions determine the identities of the adsorbent versus the scaffold. The adsorbent undergoes a cascade of wetting transitions based on the concentration of the adsorbent. To put these observations on a quantitative footing, we turned to computations to explore how adsorption depends on interaction strengths (**Fig. 3**), polymer concentration (**Ext. Data Fig. 5**), polymer length (**Fig. 4**, **Ext. Data Fig. 6**) and polymer stiffness (**Ext. Data Fig. 7**).

**Figure 5:**
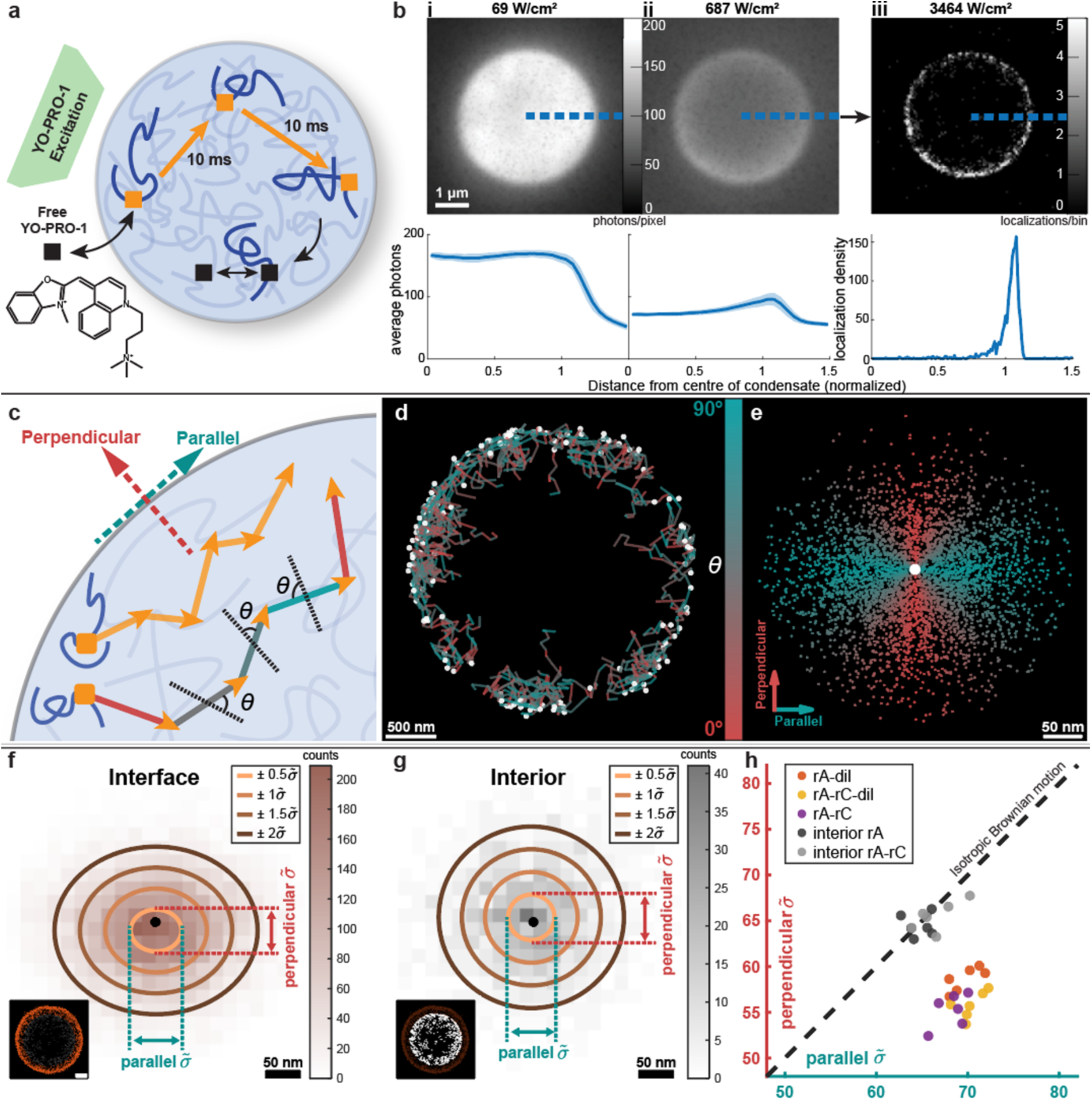
Single molecule tracking near the condensate interface shows anisotropic motion. **a**YO-PRO-1 is an RNA intercalating dye with a strong preference for binding to poly-rA over poly-rC. The dye can move into the condensates and bind to poly-rA (orange). We track the location of the dye every 10 ms until it is bleached (black). **b** The field of view within the condensate is excited at various laser strengths, bleaching increasingly more bound dye molecules until single molecules can be localized. These are seen near the interface, since the condensate is constantly bleached, and fresh dye molecules move into the condensate through the interface. **c** The trajectory in the x and y-direction is converted to the displacement parallel (aqua) or perpendicular (red) with respect to the interface. **d** Trajectories (multiple displacements) of poly-rA molecules in a poly-rA-PEG condensate. **e** Displacements of the trajectories shown in d. We fit the displacement information with 2D Gaussians for trajectories **f** near interfaces and **g** in the interior of the condensate. **h** The standard deviation in perpendicular and parallel motion for 6 condensates for trajectories near interfaces: poly-rA dense phase with dilute phase (pA-dil, orange), poly-rA dense phase with poly-rC adsorbed to the dilute phase (pA-pC-dil, yellow), and Poly-rA dense phase with Poly-rC dense phase (pA-pC, purple). Parallel motions are consistently faster than perpendicular motions. On the interior of poly-rA condensates (interior rA, dark grey) and inside of poly-rA condensates in touch with poly-rC droplets (interior rA-rC, light grey), we observe no anisotropy.

### Differential interaction strengths are the main drivers of adsorption

We modeled phase behaviors in polymer mixtures using coarse-grained, single bead per monomer representations as implemented in LaSSI^40^, which is a lattice-based simulation engine (**Figure 3**). We simulated mixtures of three distinct homopolymers labelled A, B, and C. These polymers were designed to mimic poly-rA, PEG, and poly-rC, respectively. Polymer B is parameterized to be the driver of condensate formation via heterotypic interactions with polymers A and C. The microscopic pairwise interactions between monomers of the polymers (ε_x-y_) are −2 k_B_T for most combinations, −4 k_B_T for B-C and we query the effects of varying the interaction strengths for B-A (**Figure 3a**, b, see Supplementary Methods). By maintaining equal polymer concentrations and varying ε_B-A_, we obtained three possible architectures (**Figure 3c**, d). If |ε_B-A_| = |ε_B-C_| = −4 k_B_T, then a uniform dense phase is obtained since A and C are equivalent.

For |ε_B-A_|= −4.25 k_B_T, the stronger B-A interaction increases the concentration of B and A within the dense phase and displaces polymer C to the condensate interface and dilute phase. If |ε_B-A_|= −4.5 k_B_T, polymer C is mostly excluded from the condensate interior, but is still found at the highest concentration at the interface. Adsorption is observed for the polymer C with the weaker interaction with the scaffold B. Notably, adsorption is observed in cases where polymer C partitions preferentially into the dense phase over the dilute phase (ii) and where it partitions preferentially into the dilute phase over the dense phase (iii).

For |ε_B-A_|= −4.5 k_B_T, we investigated the effect of polymer concentration on adsorption. Simulations were performed with 100 molecules of A and B, with 0, 10, 25, 50, 100, and 200 molecules of C (**Ext. Data Fig. 5 a-f**). Molecules of C adsorb preferentially to the interface, with very similar partitioning irrespective of concentration over this range. Increasing the number of A molecules or decreasing the number of B molecules reduces the number of interactions B can have with C thus increasing the dilute phase concentration of C (**Ext. Data Fig. 5 g, h**). Decreasing the number of A molecules and increasing the number of B molecules has the opposite effect, increasing the partitioning of C in the dense phase (**Ext. Data Fig. 5 i, j**). Notably, while partitioning is affected, molecules of C still adsorb, albeit weakly, to the interface, highlighting the importance of monomer interaction strength over polymer concentration.

Next, we consider the effect of polymer length on adsorption (**Ext. Data Fig. 6**). For molecules with a length of 250, |ε_B-A_|= −4 k_B_T does not result in adsorption (**Fig. 4d**). Varying the length of the molecules can change the partitioning of the molecules but it does not cause polymer C to adsorb to the interface (**Fig. 4a**). Starting in a situation where we observe adsorption for molecules with a length of 250, |ε_B-A_|= −4.5 k_B_T, varying the length of the polymers again affects partitioning. Notably, it does not prevent polymer C from adsorbing (**Fig. 4b**). Relative interaction strengths seem to be the most important variable in determining whether polymers adsorb to interfaces.

We also considered the effect of polymer stiffness on adsorption (**Ext. Data Fig. 7**). We varied molecule stiffness using a bond angle potential of the form: u(1-cos^2^8). Here, θ is the bond angle at the intersection of three consecutive beads in a chain and u quantifies the amplitude of the stiffness. For flexible polymers, we set u = 0 whereas for stiff polymers, we set u = 200. As with flexible polymers, setting |ε_B-A_|= −4 k_B_T, does not lead to adsorption even for stiff polymers (**Ext. Data Fig. 7a-f**). As far as adsorption is concerned, the results for |ε_B-A_|= −4.5 k_B_T are equivalent for stiff versus flexible polymers (**Ext. Data Fig. 7g-i**). We conclude that of the four variables considered, relative monomer interaction strength is the key determinant of condensates architecture with respect to adsorption.

### Adsorbents and scaffolds are oriented differently at the interfaces of condensates

Recently, Farag et al.,^19^ introduced an order parameter to quantify the orientation of polymers with respect to the interfaces of condensates. In their simulations, the system had only one type of polymer. Farag et al.,^19^ found that polymers at the interface of the condensate were more likely to be oriented perpendicular to the interface. This was proposed to minimize the interfacial free energy density by maximizing the number of polymers at the interface and minimizing the number of un-crosslinked stickers per polymer.

We employed a similar analysis to determine the orientations of polymers A, B and C in our simulations (**Fig. 3e**). Specifically, we determined the angle θ swept out by the condensate center-of-mass, a given polymer bead *i*, and another bead on the same polymer that is exactly 100 beads apart from bead *i*. If θ is close to 0° or 180°, which corresponds to cos^2^8 values close to 1, then the 100-bead segment is perpendicular to the interface. In contrast, if θ is close to 90°, which corresponds to cos^2^8 values close to 0, then the 100-bead segment is essentially parallel to the interface. In the case of a polymer that has a random orientation, cos^2^8 = 0.33 (**Ext. Data Fig. 5k**).

When |ε_B-A_| = |ε_B-C_|= −4 k_B_T, all polymers have random orientations, except at the interface, where there is a bias toward perpendicular orientations. This matches results by Farag et al.^19^. We also computed the orientational preferences in the case of adsorption realized for |ε_B-A_|= −4.25 or −4.5 k_B_T. In the dense and dilute phases, orientations are random. However, at the interface, polymer A and B have an enhanced perpendicular orientation. At the highest concentration of polymer C, approximately 30 l.u. from the center of the condensate, we observe that polymer C has a more parallel orientation. These results suggest that polymers that behave as scaffolds, namely polymers A and B, minimize the interfacial free energy density by adopting more orthogonal orientations at the interface. In contrast, the homopolymer C effectively minimizes its interfacial free energy density by showing adsorbent-like behavior. Notably, the ability to adsorb and obtain this orientation is not intrinsic because polymer C can form condensates on its own in the appropriate concentration regimes. Instead, condensate formation by the scaffold molecules causes adsorption if the relative interaction strengths are such that |ε_B-A_| > |ε_B-C_|. Interfacial adsorption maximizes interactions with the condensate interface while minimizing the disruption of favourable A-B interactions that provide the cohesive interactions for condensate formation and stabilization.

### Observations of anisotropic motions near condensate interfaces

Given predictions of specific orientational preferences of scaffolds versus adsorbents at the interface, we asked if molecules near the interface also display anisotropic motions. We tracked the movement of single poly-rA molecules (blue) in condensates near the interfaces (**Fig. 5a**). YO-PRO-1 is an RNA binding dye which can diffuse into a poly-rA condensate and intercalate with the RNA. Once bound to RNA, we excite the dye (yellow squares). Over time and depending on the laser power, some of the dyes can be bleached (black square). At low laser power, the poly-rA-rich phase is bright, while the dilute phase is dim (i, **Fig. 5b**). Increasing the laser power results in significant bleaching and the middle of the poly-rA-rich phase becomes dark (ii, **Fig. 5b**). At very high laser power, we observe mostly poly-rA molecules near the interface, due to the combination of constant bleaching and new dye molecules entering the condensate via the interface (iii, **Fig. 5b**). The high laser power allows us to collect single-molecule super-resolution images every 10 ms and eliminates signals further away from the interface. We thus obtain trajectories of the poly-rA molecules, which we convert from displacements in x and y-direction to displacements parallel or perpendicular to the interface for each 10 ms step (**Fig. 5c, d**). Plotting each displacement starting from the same location, we gain an overview of movement of molecules (**Fig. 5e**). We split these data into displacements within 300 nm of the interface (**Fig. 5f**) and further away from the interface, labelled “Interior” (**Fig. 5g**) (**Ext. Data Fig. 8**). We then fit over 10^4^ displacements with a 2D Gaussian obtaining the mean and standard deviation (ellipses) traversed in the parallel and perpendicular directions from the starting location (black dot). We find that near the interface, movements parallel to the interface are significantly faster than movements perpendicular to the interface. This effect disappears for movements in the interior of the dense phase. **Fig. 5h** shows the standard deviation of poly-rA molecules movement of six condensates near three types of interfaces: poly-rA dense phase with dilute phase (pA-dil, orange), poly-rA dense phase with poly-rC adsorbed to the dilute phase (pA-pC-dil, yellow), and poly-rA dense phase with poly-rC dense phase (pA-pC, purple). These data points are off the diagonal, supporting the conclusion that motions of the dyes parallel to the interface are consistently faster than motions of the dye that are perpendicular to the interface. We do not observe anisotropies of motions within the interior of poly-rA-rich phases (interior rA, dark grey) or inside poly-rA-rich phases that are in contact with poly-rC-rich phases (interior rA-rC, light grey).

Our findings regarding the anisotropies of motions at interfaces are in accord with predictions from theory ^41^. For a sphere near a liquid-liquid interface, a higher drag in the perpendicular direction, which translates to lower speeds in this direction when compared to the parallel direction was predicted as being a consequence of the incompressibility of a fluid-fluid interface ^41–43^. Our measurements appear to be the first study confirming this prediction for anisotropy in molecular motions at interfaces.

## Discussion

Our experiments and simulations paint a detailed picture of how molecules behave at interfaces between coexisting dense and dilute phases formed by ternary mixtures of RNA molecules and PEG. We report that the identities of adsorbents versus scaffolds are governed by the relative monomer interaction strengths among different molecules. The conventional wisdom in the condensate field is that molecules can be easily classified as being scaffolds or clients. However, our work shows that clients, defined as a molecules of lower valence or weaker interactions are not just passive molecules that partition into dense phases formed by scaffolds. Instead, in a system comprising molecules A, B, and C where A and C interact favorably with B, we obtain the following results: If the A-B interactions are stronger than the B-C interactions, then the C molecules not only partition into the A-B-rich phase, but they also adsorb onto the interface of the A-B-rich phase. Adsorption appears to be the consequence of a balance whereby the C molecules cannot outcompete the stronger A-B interactions, and they can gain access to B molecules at the interface that are not engaged in interactions with A molecules. Increasing the concentration of the C molecules, such that the threshold for phase separation with B molecules is crossed, leads to a cascade of wetting transitions that eventually lead to A-rich, C-rich, and dilute phases coexisting with one another. While polymer concentration, length and stiffness can affect partitioning into the dense or dilute phase significantly, our computations show that the central determinant of adsorption and condensate architecture is the relative monomer interaction strength.

In contrast to coexisting dilute or dense phases, molecules at interfaces have non-random orientational preferences with respect to the planes of interfaces. The drivers of phase separation namely the scaffolds have a more perpendicular orientation to the interface whereas the weaker interactors, which are adsorbents prefer more parallel orientations. Our study identifies ways to tune condensate architectures and interfacial structures by modulating the relative strengths of scaffold versus adsorbent interactions. The structured interface also influences the motion of molecules, causing movement in the parallel direction to be faster than in the perpendicular direction.

Our observations, which were derived using generic, biocompatible, and biologically relevant homopolymers, help explain the phenomenology of adsorption that has been reported in cells ^32,44^. Further, since multiphase condensates create internal interfaces ^14,15^, we propose that the internal interfaces between dense phases might provide additional spatial and temporal control over biochemical interactions. Specifically, interfaces between dense phases, as well as interfaces between coexisting dilute and dense phases, are likely to provide distinct molecular microenvironments for molecules to accumulate and interact. Adsorption and wetting transitions at interfaces are also relevant for the design of condensates with tunable catalytic functions, given observations from the microdroplet literature which show that reaction efficiencies can be enhanced by orders of magnitude at the interfaces of coexisting phases ^45^. Additionally, recent studies have shown that condensates are defined by interphase electric potentials ^46^, that help set up interfacial electric fields ^47^. These fields enable the catalysis of reactions such as hydrolysis of ATP^48^. Furthermore, in spatially organized condensates, there can be internal and external interfaces that can be engineered to minimize off-target reactions in biochemical reactions involving multi-enzyme pathways ^45^.

Given recent advances and the growing focus on interfaces, our work shows how rules can be extracted and deployed in future designs of condensates with bespoke interfacial structure and dynamics. The mechanisms of adsorption and wetting are distinct from the site-specific effects of ligands that are known to modulate driving forces for phase separation through preferential binding across phase boundaries ^49–52^. While preferential binding of ligands ^53^ can either compete against or add to extant interactions, thereby destabilizing or stabilizing condensates, adsorption of a ternary component has little effect on the interactions within a condensate ^54^. Overall, our results highlight key physical principles that govern the formation of spatially heterogeneous condensates of the type observed in live cells. Additional consideration of site- or chemistry-specific associative interactions are likely to provide further tunability to the spatial organization of macromolecules within and around condensates.

## Supplementary Information

Detailed description of the design of the experiments, reagents used, analysis pipelines and computational methods are available in the supplementary information document.

## Acknowledgements

This work was funded by the Royall Scholarship (N.A.E.), the Center for Biomolecular Condensates at Washington University in St. Louis (N.A.E., R.V.P.), the European Union’s Horizon 2020 research and innovation program under the Marie Skłodowska-Curie grant MicroREvolution (agreement no. 101023060; T.S.), the Winston Churchill Foundation of the United States (T.J.W.), the Harding Distinguished Postgraduate Scholar Program (T.J.W.), Global Research Technologies Novo Nordisk A/S (H.A., T.P.J.K.), the US National Institutes of Health (R.V.P), the US Air Force Office of Scientific Research (R.V.P), the European Research Council under the European Union’s Seventh Framework Program (FP7/2007-2013) through the ERC grants PhysProt (agreement no. 337969; T.P.J.K.) and the Newman Foundation (T.P.J.K., T.S.).

## Author Contributions

N.A.E., T.S. and T.P.J.K. conceived the study. N.A.E., M.F., Y. Q., D.Q., T.S., T. W., T.J.W., H.A., T.J.K., D.A.W., M.L., T.P.J.K. and R.V.P. conducted investigations and interpreted the results. T.P.J.K. and R.V.P. acquired funding. N.A.E., M.F., T.P.J.K. and R.V.P. wrote the original drafts, all authors reviewed and edited the paper.

## Conflict of interest

RVP is a member of the scientific advisory board of and shareholder in Dewpoint Therapeutics Inc. TPJK is co-founder of Fluidic Analytics, Wren Therapeutics, Xampla, and Transition Bio. These associations did not influence the work reported here. All other authors have no conflicts to disclose.

## Additional information

Data generated during the study are available on request from the corresponding authors.

## Extended Data Figures

**Extended Data Fig. 1.**
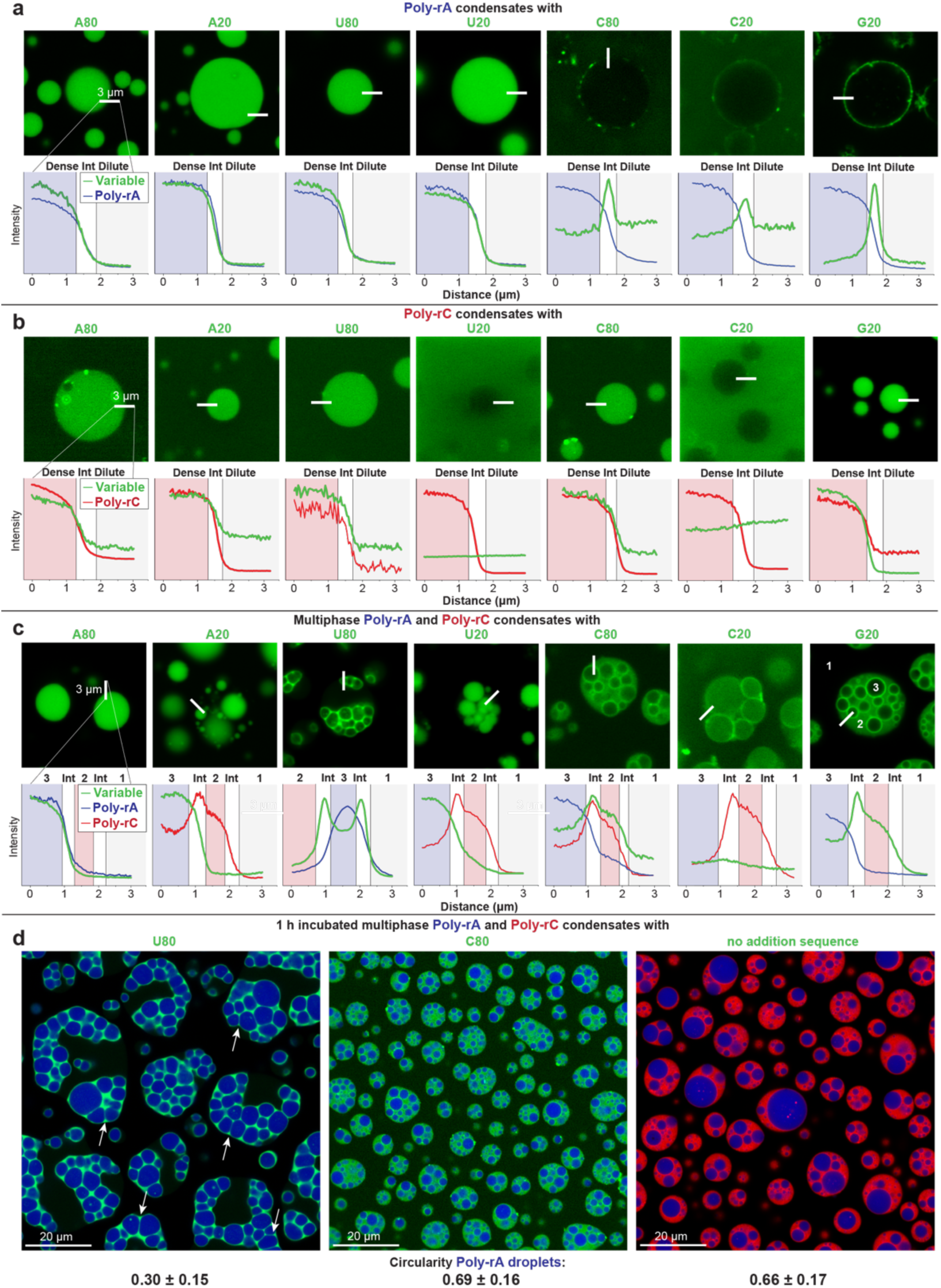
Phase separation and adsorption of poly-rA and/or poly-rC solutions. rA_80_, rA_20_, rU_80_, rU_20_, rC_80_, rC_20_, rG_20_ were added to condensates formed by: **a** poly-rA, **b** poly-rC and **c** poly-rA and poly-rC. **a** For poly-rA condensates, rC_80_, rC_20_ and rG_20_ adsorb onto the interface. **b** For poly-rC condensates, the short RNA sequences do not show significant adsorption onto the interface. In panels a and b, Int refers to the interface. **c** For poly-rA and poly-rC condensates, we observe strong adsorption of rU_80_ to the interface. Similar results were obtained for rC_80_, rC_20_ and rG_20_. In these experiments, we used poly-rX (1g/L), rX_80/20_ (0.1 g/L), 750 mM NaCl, 5 w/w% PEG, 50 mM Tris pH=7.5. **d** Adsorption onto poly-rA and/or poly-rC condensates can prevent fusion. To the poly-rA-poly-rC multiphase system, we added **a** rU_80_, **b** rC_80_ or **c** nothing. rU_80_ is complementary to poly-rA. The rU-rich sequence adsorbs strongly onto the interface and prevents the poly-rA droplets from becoming spherical. We observed higher sphericity for rC_80_ or no additional component. All scale bars in the confocal images represent 3 μm.

**Extended Data Fig. 2.**
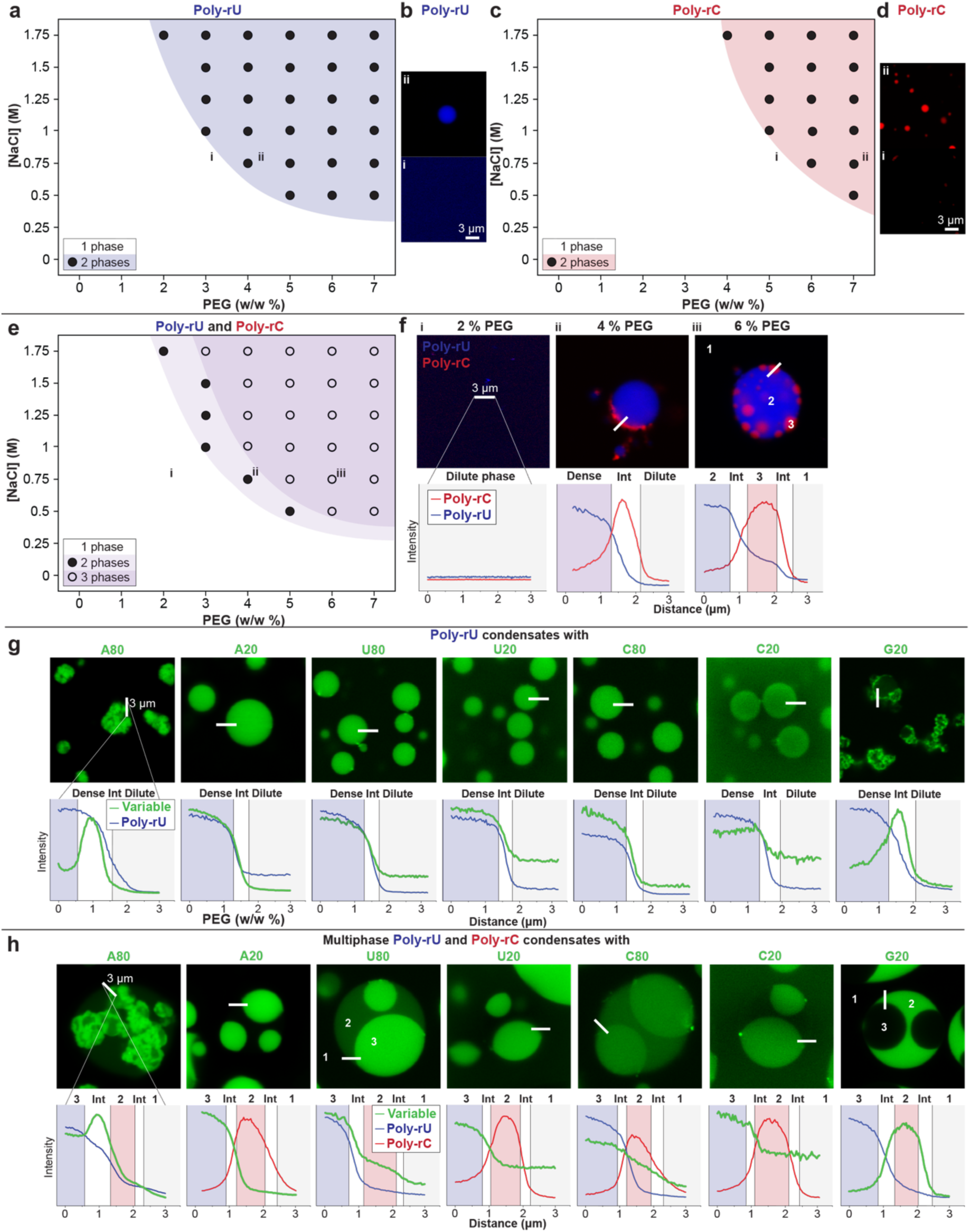
Phase separation and adsorption of poly-rU and poly-rC. **a**Poly-rU (1 g/L) can phase separate in the presence of sufficient [NaCl] and/or [PEG], **b** forming condensates. **c** Poly-rC (1 g/L) phase separation requires more [NaCl] and/or [PEG] **d** to form condensates. **e** The phase diagram of a mixture of poly-rA (1 g/L) and poly-rC (1 g/L). At low [PEG] (i of **f**) one homogenous phase is obtained. Increasing [PEG], poly-rU forms condensates and some poly-rC is recruited into it (ii of **f**). Notably, poly-rC does not significantly adsorb to the interface. Increasing [PEG] leads to poly-rU phase separation (iii of **f**). rA_80_, rA_20_, rU_80_, rU_20_, rC_80_, rC_20_, rG_20_ is added to **g** poly-rU and **h** poly-rC and poly-rU condensates. **g** For poly-rU condensates, rA_80_ and rG_20_ forms aggregate-like structures, which stick to the interface of the condensate, functioning like Pickering emulsions. **h** For poly-rU and poly-rC condensates, we observe strong adsorption of U80 to the interface to the interface. The rest of the sequences prefer one of the dense phases. Poly-rX (1g/L), rX_80/20_ (0.1 g/L), 750 mM NaCl, 5 w/w% PEG, 50 mM Tris pH=7.5. All scale bars in the confocal images represent 3 μm.

**Extended Data Fig. 3.**
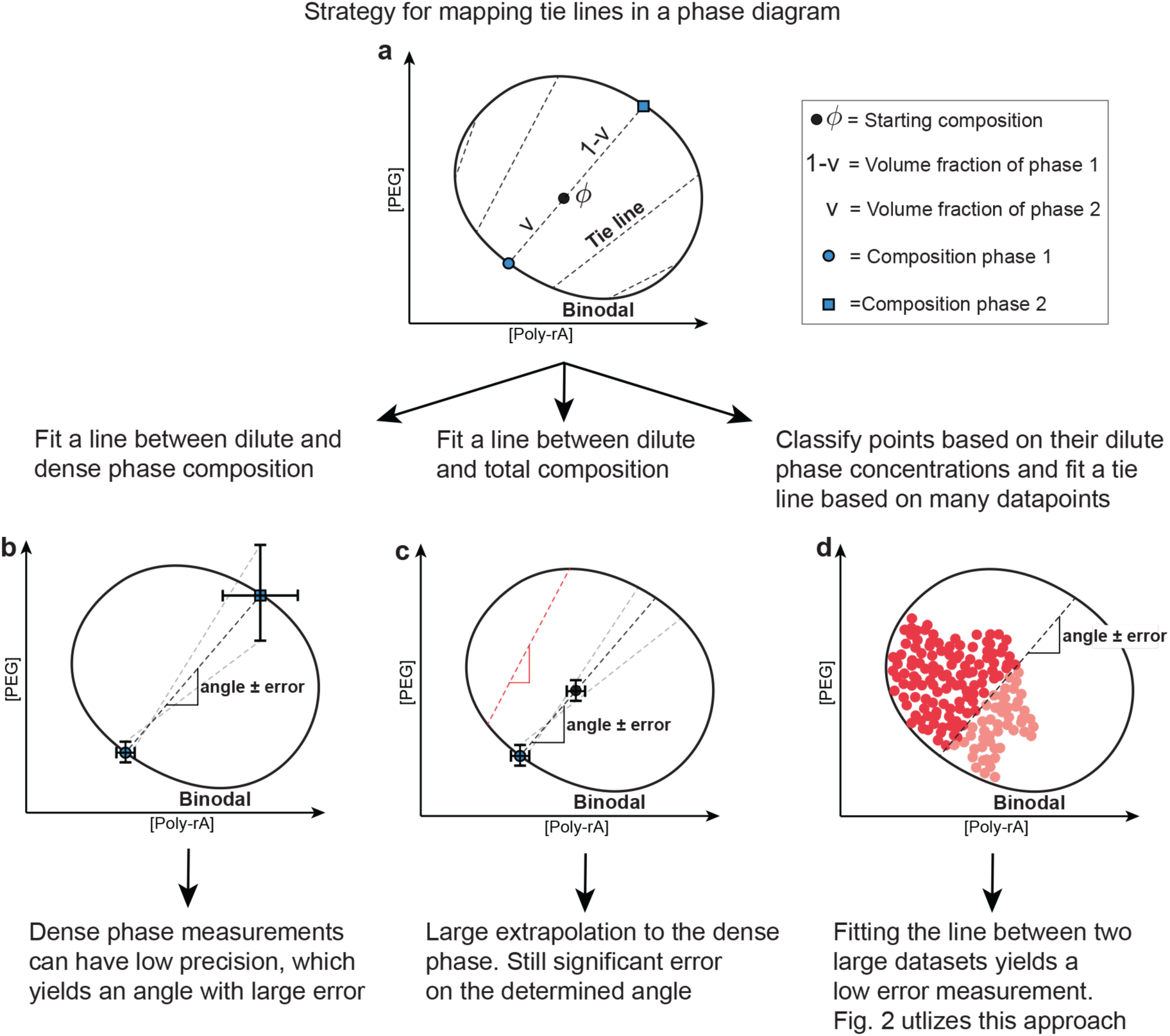
Strategies for measuring tie line proxies. **a**An example phase diagram for the molecules PEG and poly-rA. Here, we draw the tie line (dashed) that connects the dilute phase (phase 1, blue circle) and the dense phase (phase 2, dense phase or condensate). Note that the tie lines are not parallel. Considering different options for measuring tie lines, we find that **b** measuring the dense phase concentrations involves significant errors and is difficult to automate, and **c** that fitting between the total and dilute phase is easier to automate, but also associated with significant errors due to the proximity of the datapoints. **d** The overall procedure is as follows: Fluorescence intensities converted to total concentrations are used to place the datapoints in the phase diagram, defining the input or total concentrations. Phase separated (red) datapoints and mixed datapoints (blue) are used to map the phase boundary. To determine proxies of tie lines, we determine the concentration of RNA in the coexisting dilute phase for all the points on the plane that correspond to the input (i.e., total) concentration. All the total compositions that correspond to a specific value for the concentration of RNA in the coexisting dilute phase fall on a single tie line. To take advantage of the thousands of points that can be scanned and coexisting dilute phase concentrations that can be measured, we use the fact that tie lines cannot cross, and segment data into two sets of points that must lie above and below a tie line proxy. This segmenting procedure, based on a support vector machine classifier, is used to obtain proxies for tie lines that best separate adjacent sets of data into two distinct sets. Repeating the procedure across the set of points we gather, we can locate a series of lines that segment the data, which involves knowledge of the input concentrations (or total concentrations) and measurements of concentrations in the coexisting dilute phase. The procedure is used to find ∼10 tie line proxies per phase diagram.

**Extended Data Figure 4:**
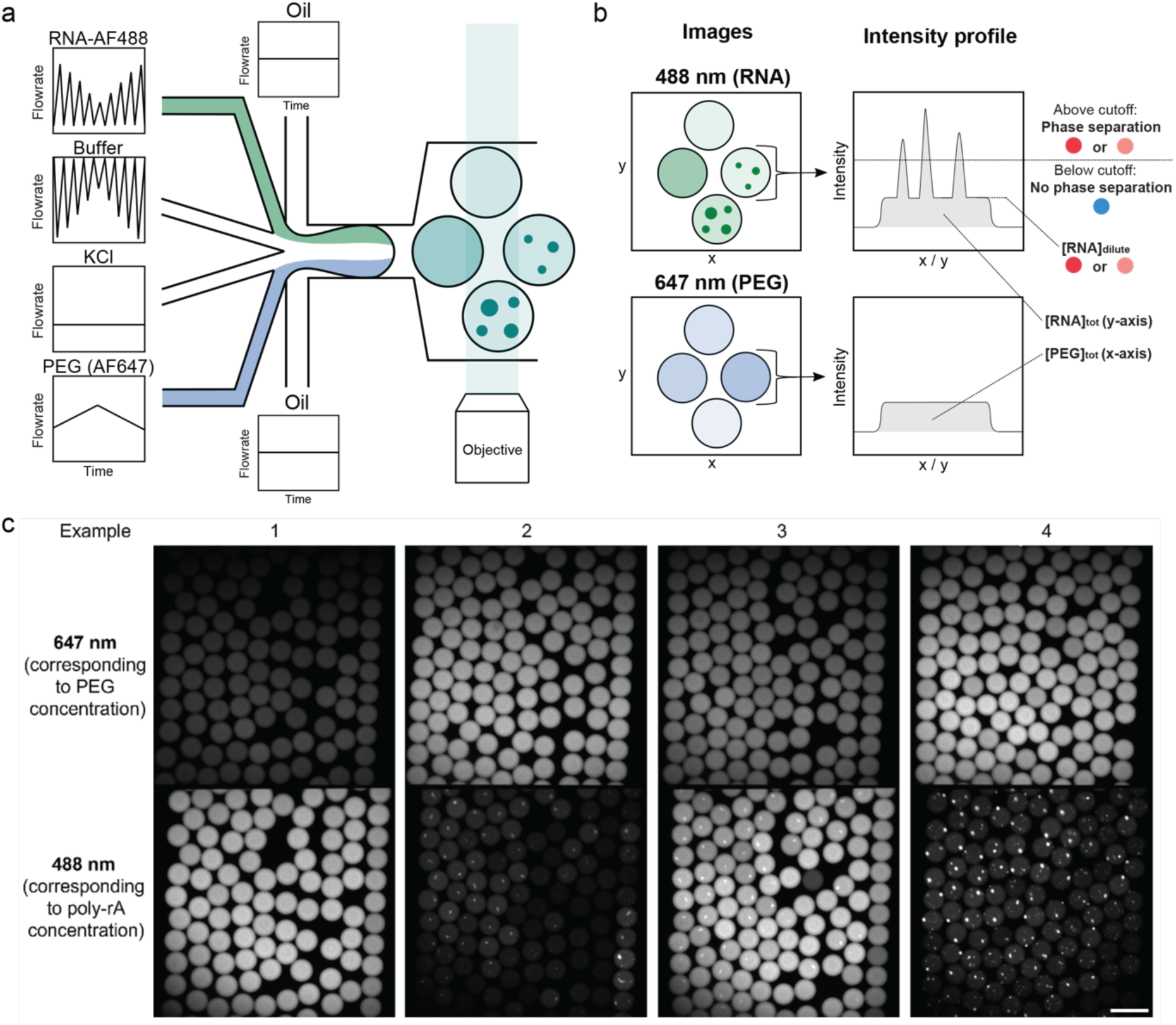
Measurements of phase boundaries using microfluidics. **a.** Schematic describing the microfluidic platform where droplets of approximately 100 pL are prepared. Flow rates of RNA, buffer and PEG solutions are varied to create samples with different compositions. **b.** Samples are imaged at 488 and 647 nm, creating pictures showing the location of the RNA and the barcode dye for PEG concentration. From the total 488 intensity in samples, we can find the total RNA concentration. Additionally, we observe if samples are phase separated (condensates can be seen as bright specs) or mixed (homogeneously distributed dye). In the case of phase separated droplets, we can additionally determine the dilute phase concentration. The intensity of the 647 dye indicates the concentration of PEG. Calibrations to convert intensity to concentration and to correct for illumination differences between locations in the pictures were performed. The information obtained from these samples is used to construct Main text Figure 2d and e. **c** Example raw images of the water in oil samples. Note that the same droplets are shown twice, imaged at different wavelengths. Samples in the same image can have different compositions or intensities. The scale bar represents 100 μm.

**Extended Data Figure 5:**
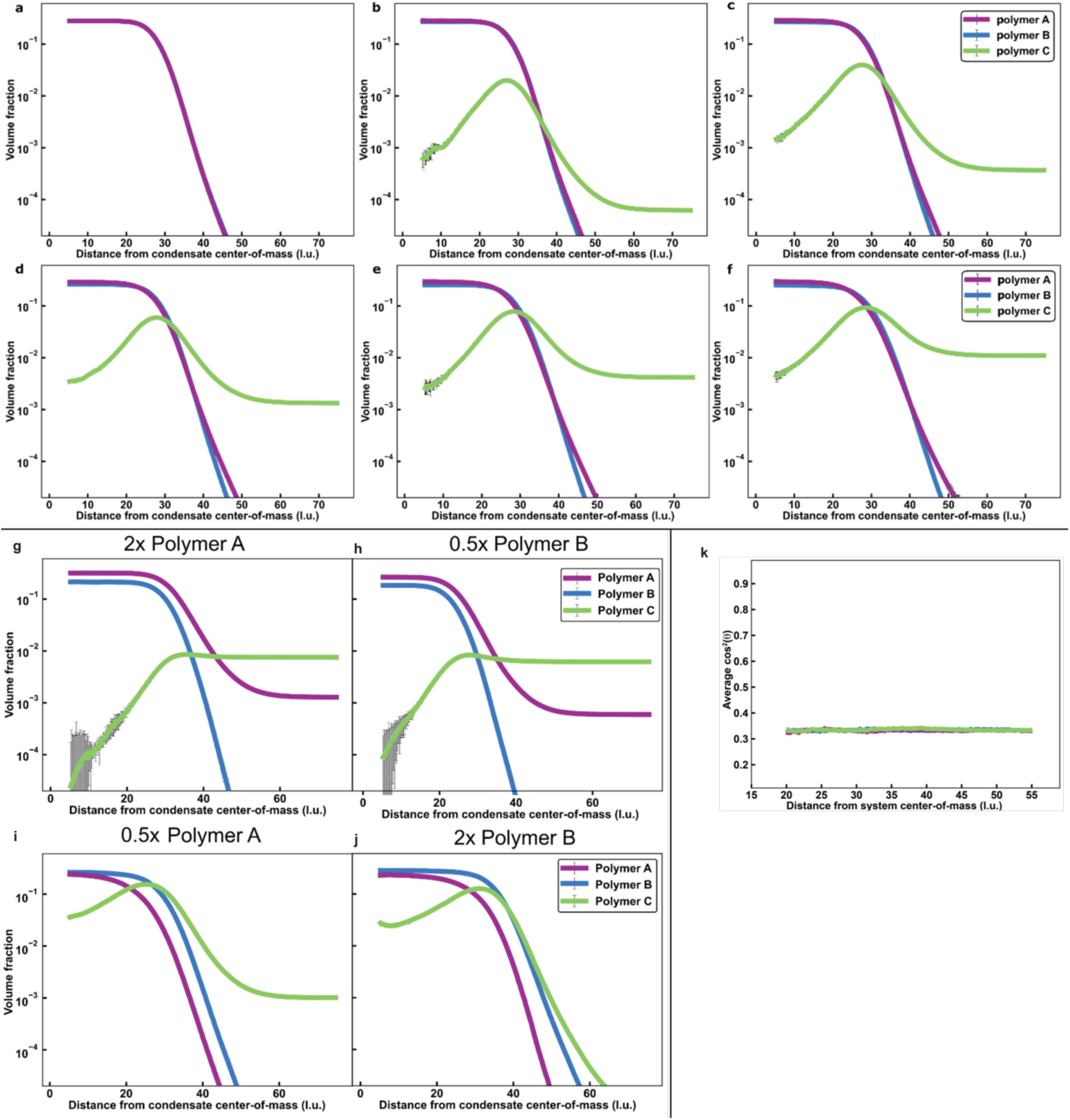
Radial density profiles from LaSSI simulations probing the effects of varying the numbers of C polymers and / or the ratio of A-to-B polymers. For |ε_B-A_|= −4.5 k_B_T (Fig. 3), we determine the radial density profiles with 100 molecules of polymers A and B, respectively. We investigate the effects of varying the numbers of polymer C from **a** 0, **b** 10, **c** 25, **d** 50, **e** 100, and **f** 200. We also investigate samples containing 100 molecules of C. Additionally, we probe the effects of there being different numbers of A and B polymers: **g** 200, 100 ;**h** 100, 50 ; **i** 50, 100 ; **j** 100, 200. Adsorption of polymer C is largely independent of the concentration of C, within the ranges that were investigated. Adsorption of polymer C is promoted by increasing the amount of polymer B or decreasing the amount of polymer A. **k** In the one phase regime, the orientations of the molecules are random, thus cos^2^8 = 0.33 at all distances for each polymer species. The relative interaction energy between polymer A and polymer B beads is 0.67. Error bars indicate the standard errors about the mean across three replicates.

**Extended data Fig 6.**
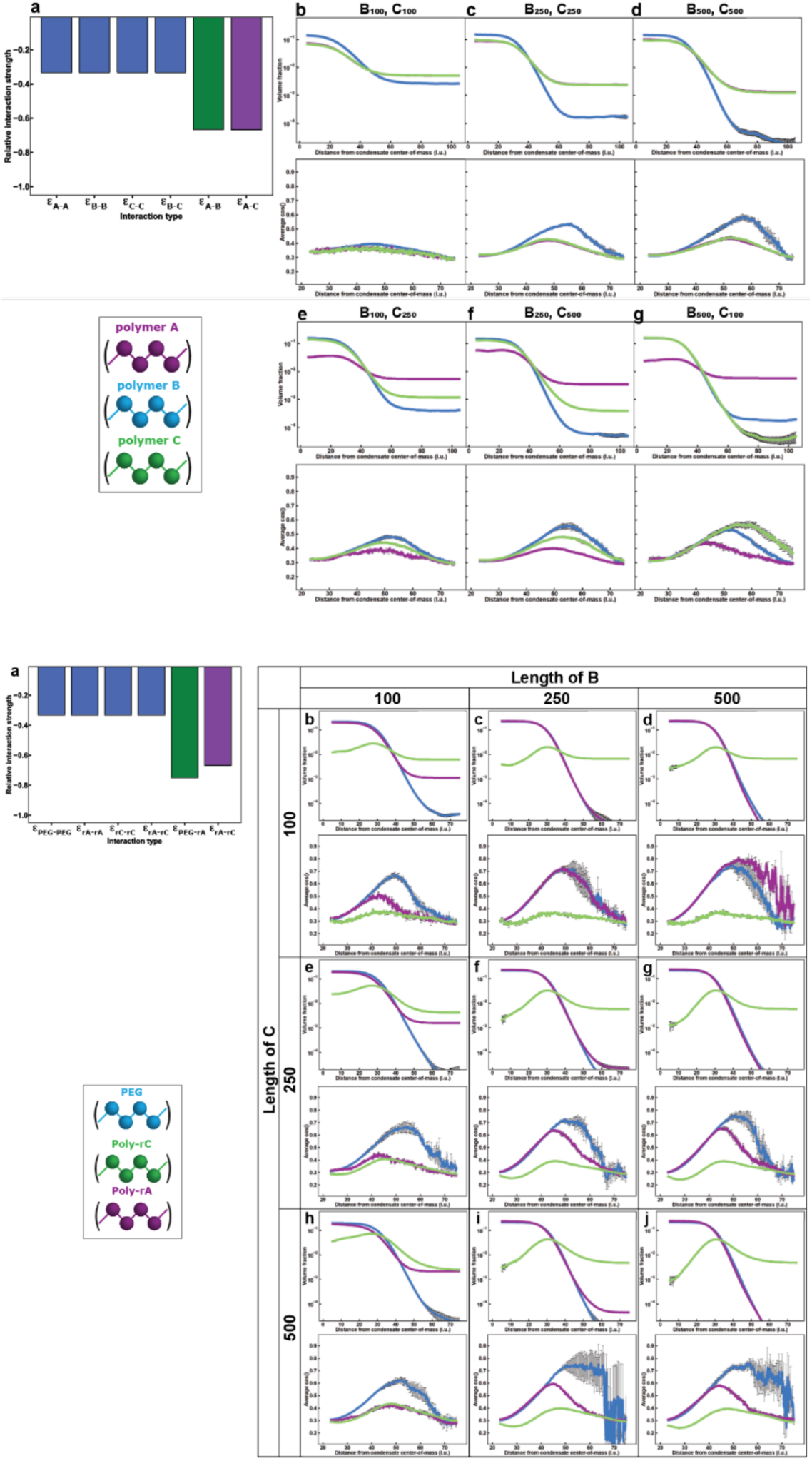
LaSSI simulations designed to mimic the effects of varying the lengths of poly-rA and poly-rC and PEG. Two sets of simulations were performed to determine the effect of polymer length on adsorption and orientation. We plot the radial density profiles in simulations and average cos^2^ 8 values, which indicate the average orientation of molecules with respect to the center of the condensate **a** For |ε_B-A_|= −4 k_B_T, we perform simulations for the following lengths of polymers that mimic poly-rA and poly-rC of: **b** 100, 100; **c** 250, 250; **d** 500, 500; **e** 100, 250; **f** 250, 500; **g** 500,100. **a** For |ε_B-A_|= −4.5 k_B_T, we perform simulations for different lengths of poly-rA and poly-rC of: **b** 100, 100; **c** 100, 250; **d** 100,500; **e** 250, 100; **f** 250, 250 (similar to Fig. 3); **g** 250,100; **h** 500, 100; **i** 500, 250; **j** 500,500. The results show that the polymer length influences the ability of a molecule to act as a scaffold, and thus influences the partitioning into the dense phase. However, the length has very little effect on whether a molecule will act as an adsorbent. Error bars indicate the standard errors about the mean across three replicates.

**Extended Fig 7.**
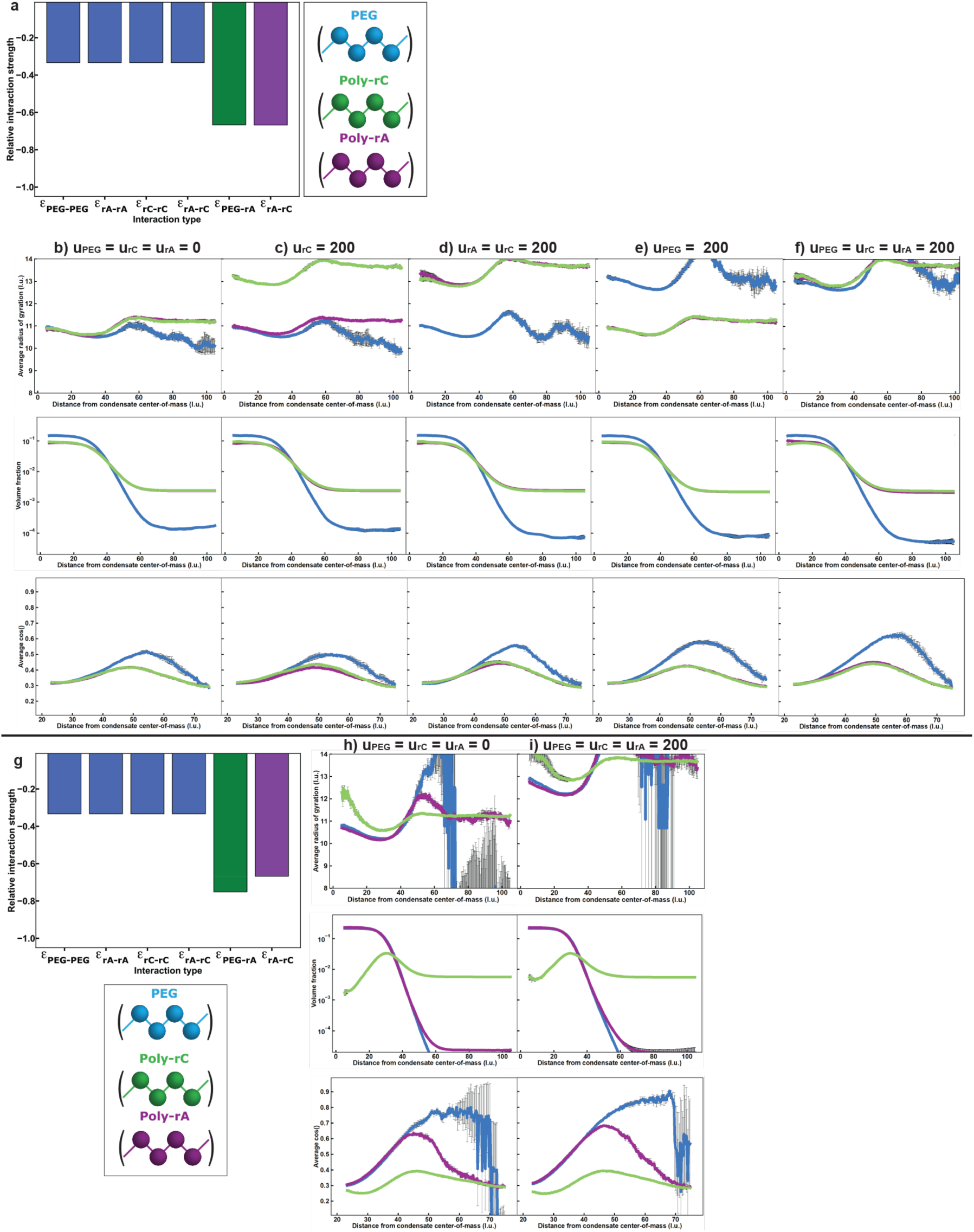
LaSSI simulations varying molecule stiffness. We plot the average radius of gyration, the radial density profiles and average cos^2^ 8 values. We vary molecule stiffness, using the potential u[1-cos^2^(θ)] where θ is the angle swept out by three consecutive beads in a chain and u is a stiffness parameter. **a** For |ε_B-A_|= - 4 k_B_T, we set **b** u(PEG)=u(rC)=u(rA)=0; **c** u(PEG)=u(rA)=0 u(rC)=200; **d** u(PEG)=0 u(rC)=u(rA)=200; **e** u(PEG)=200 u(rC)=u(rA)=0; **f** u(PEG)=u(rC)=u(rA)=200 **g** u(PEG)=u(rC)=u(rA)=0; **h** u(PEG)=u(rC)=u(rA)=200. From these simulations we conclude that the radius of gyration increases significantly when the stiffness is set to 200 instead of 0. However, the location of the molecules, and extent of adsorption is minimally affected when compared to the fully flexible case. While the extent of adsorption is minimally affected, the orientation at the interface is slightly more perpendicular for stiff molecules over more flexible ones. Error bars indicate the standard errors about the mean across three replicates.

**Extended data Fig. 8.**
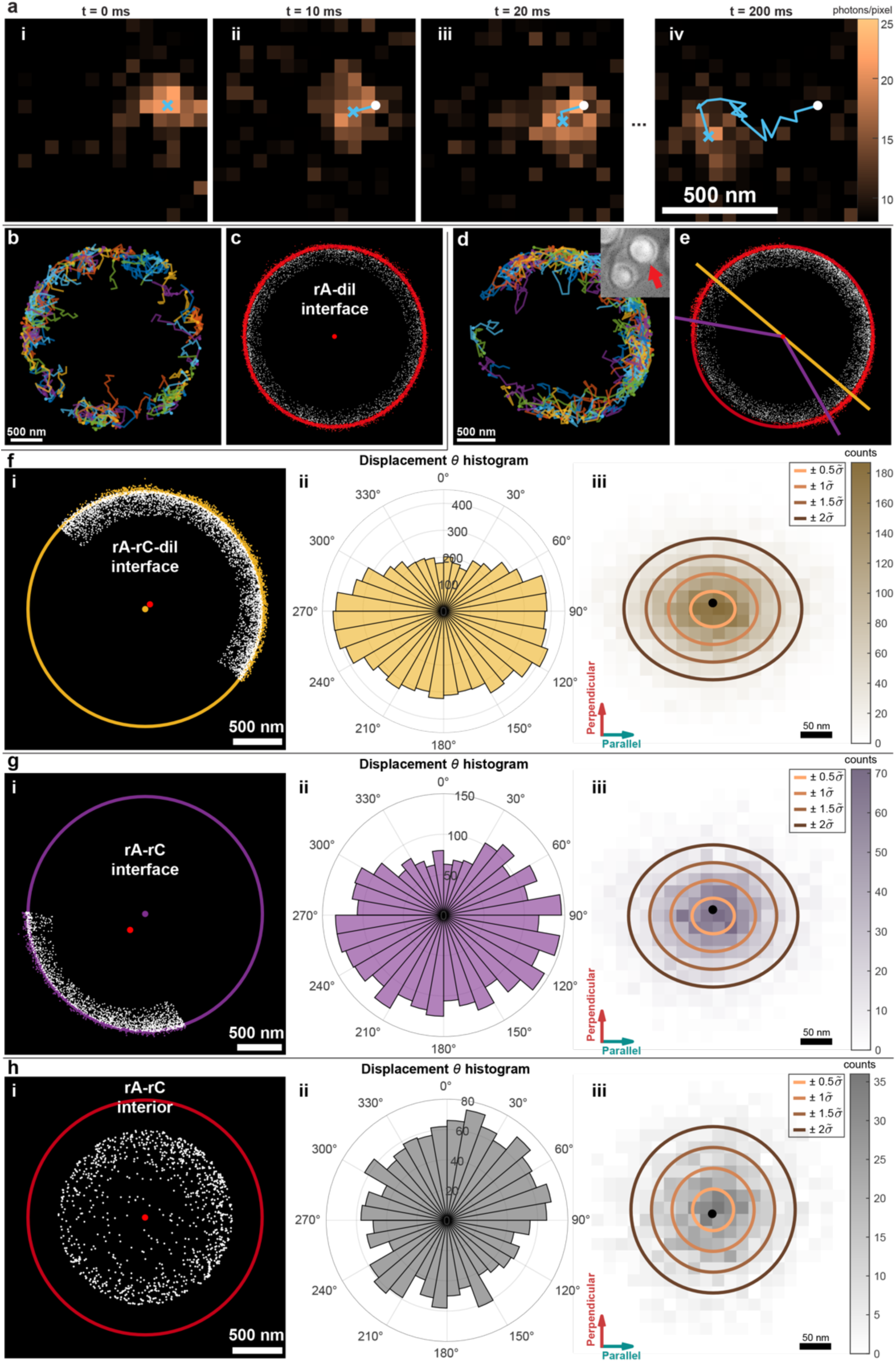
Condensate imaging, localizing single molecules and displacement calculations. For poly-rA condensates and their interface with the dilute phase, we show **a** the movement of a molecule over 200 ms, **b** all trajectories of 60 ms or longer, **c** all starting points of trajectories with the fitted interface as a reference point to determine **d** the displacements perpendicular (up/down) and parallel (left/right). **e** For a poly-rA condensate with an interface with the dilute phase (rA-rC-dil) and an interface with the poly-rC dense phase (rA-rC), we divide the data based on the closest interface. The data for **f** the rA-rC-dil interface and the **g** rA-rC interface are fitted with separate reference circles to account for different contact angles of the phases. **h** We also study displacements further away from the interface (>300 nm). We obtain the perpendicular and parallel displacement and add this data to Fig. 5h.

