## Supplemental Materials for "Differential interactions determine anisotropies at interfaces of RNA-based biomolecular condensates"

Contains:

- Supplementary materials and methods
- Supplementary references

### Supplementary Materials and Methods

#### Materials

Poly-rA (MW 700-3500 kDa, 10108626001), PEG (average MW ~20,000), Poly-rC (average MW ~118 kDa, P4903), HEPES, NaCl, KCl, ethanol and mPEG(5k)silane were obtained from Sigma Aldrich. Alexa Fluor™ 647 Carboxylic Acid, Alexa Fluor™ 488 Carboxylic Acid and RNAase free water were obtained from Thermo Fisher. PEG(20k)-AF647 was purchased from Nanocs. Sylgard 184 Elastomer base and curing agent were purchased from Dow Corning Corporation. 18×18 mm glass slides were purchased from Academy. 24×60 mm No.1.5 glass slides were purchased from DWK Life Sciences. $(T)_{20}$-AF488 and $(C)_{20}$-AF546 were purchased from GenScript.

#### Fabrication of imaging wells for confocal microscopy

PDMS slabs of 3 mm in height were fabricated by mixing Sylgard 184 Elastomer base and curing agent at a 10:1 ratio and heating at 65 °C for 1.5 hours. Holes of 5 mm in diameter were punched in PDMS slabs of ~3 mm height using biopsy puncher. These slabs were plasma bonded to 24×60 mm No.1.5 cover glass slides to create wells. Both the wells and 18×18 mm glass slides used to seal the top of wells were treated with PEG-silane, using a method based on previously reports[3,4]. Briefly, the treatment solution was prepared by mixing 10 mg of PEG(5000)silane with 20 µL of glacial acetic acid and 1 mL of ethanol. Wells and glass slides were treated by placing them in solution for 1 hour at 65 °C and afterwards washing them thoroughly with water. Treated devices are not used after more than 3 weeks since the treatment.

**Condensate preparation**

All condensates are prepared and imaged at room temperature. Condensates of poly-rA and or poly-rC with PEG were prepared by mixing stock solutions in an Eppendorf tube in the following order: PEG, buffer, RNA, salt. All solutions were prepared in RNAase free water. The poly-rA and poly-rC concentrations were determined with a nanodrop machine by measuring the absorbance at 260 nm. Poly-rA and poly-rC were fluorescently labelled by incubation of 0.5 mol-% of $(T)_{20}$-AF488 or $(C)_{20}$-AF546 for 3 minutes at 80 °C. The sample was placed in an imaging well and sealed airtight.

**Confocal imaging**

A Leica Stellaris 5 confocal microscope (confocal fluorescence imaging) (white light laser) microscope equipped with a 63× oil immersion Leica 1.4 NA was used for imaging. Further analysis of pictures was performed using Fiji.

**Microfluidic device design and fabrication**

Standard lithography processes were used to fabricate the microfluidic devices.[2,5] AutoCAD (AutoDesk) software was used to design the devices, which were printed on a photomask (Micro Lithography). A 45 µm layer of SU8-3050 photoresist (Microchem) was spun on a silicon wafer and heated at 95 °C for 15 minutes. The photomask was placed on top, and the pattern was placed onto the wafer using exposure to UV light and heating at 95 °C for 15 minutes. Propylene glycol methyl ether acetate (PGMEA) was used to remove excess unreacted SU8-3050. The wafer was dried and a profilometer was used to confirm the height of the device. PDMS at 1:10 curing agent:base ratio was placed on top of the mould and heated at 65 °C for 1.5 hours. The PDMS was cleaned, holes were punched into inlets and outlets, the PDMS substrate was then bonded to glass slides using a plasma oven (30 s, 40% power, Femto, Diener Electronics). The interior of the device was treated with 1% trichloro(1H,1H,2H,2H-perfluorooctyl)silane (Sigma) in HFE-7500 (Fluorochem), before being dried and heated at 95°C for 1 min.

**Mapping phase diagrams and constructing tie-lines**

Aqueous droplets in oil of an average volume of 100 picolitre were created at approximately 100 droplets per second using a microfluidics setup as shown schematically in extended data figure 4. Specifically, a solution of fluorescently labelled $10^4$ ng/µL poly-rA or poly-rC was mixed with a solution of fluorescently labelled 20 w/w % PEG and a solution with buffer at varying rates while a constant amount of KCl solution was added, all of which were being encapsulated in fluorinated oil (HFE-7500 Fluorochem with 1 v/v % fluorosurfactant), flowing at constant speed. To flow the solutions into the microfluidic device, syringes (Hamilton 1710) and syringe pumps (neMESYS modules, Cetoni) were used. Syringes were connected to the microfluidic device using tubing (PTFE, 0.012"ID × 0.030"OD, Cole-Parmer). The 100 picolitre droplets were created by a total aqueous flow rate of 60 µL/h and the oil flow rate of 150 µL/h. After a 5-minute incubation period, the aqueous droplets in oil were imaged simultaneously at the wavelength corresponding to poly-rX (488 or 546 nm) and PEG (647 nm). These images were analysed using the Python program[1] to determine the concentrations of these compounds and the presence or absence of condensates. Additionally, the dilute phase concentration of poly-rX was determined by calibrating the intensity of 488 or 546 in an area absent of condensates to concentration poly-rX concentration. Each droplet gives information for one datapoint in figures 2d or e. The boundaries were determined using fitting that is based on support vector machine-based methods. The tie-lines were fitted between the "condensates" datapoints by grouping them based on their poly-rX dilute phase concentration.

**Coarse-grained simulations**

Simulations were performed using LaSSI, a lattice-based Monte Carlo engine[6]. Monte Carlo moves are accepted or rejected based on the Metropolis-Hastings criterion so that the probability of accepting a move is equal to min(1, exp(−βΔ*E*)), where β = 1 / *kT*. Here, *kT* is the simulation temperature (set to 55) and Δ*E* is the change in total system energy associated with the attempted move. Total system energies were calculated using a nearest neighbour model and the following interaction energies: $e_{A\text{-}A}$ = $\varepsilon_{B\text{-}B}$ = $\varepsilon_{C\text{-}C}$ = $\varepsilon_{A\text{-}C}$ = –2; $\varepsilon_{B\text{-}C}$ = –4; $\varepsilon_{B\text{-}A}$ was -4, -4.25, or -4.5. Unless stated otherwise, each simulation involves $10^2$ distinct polymers of each type comprised of 250 beads each in a cubic lattice with length 150 lattice units. The simulations were performed for $5\times10^{10}$ Monte Carlo moves and were allowed to equilibrate such that the chains formed a single condensate with a coexisting dilute phase before any analysis was performed. Three independent simulations were performed at each condition, and the results shown are aggregates over all replicates. Radial densities of each polymer type were normalized using the exact number of lattice sites in each radial shell. The condensate centre-of-mass was taken to be the centre-of-mass of the single largest network of interacting chains. The chain orientation analysis in figure 3 was introduced previously[6]. Here, we perform the analysis as follows: (1) for a given chain in the system, find all pairs of beads (*i*, *j*) that are exactly 100 beads apart along the chain. (2) For each pair, draw a line segment from the condensate centre-of-mass to bead *i*. (3) Draw a line segment from bead *i* to bead *j*. (4) Determine the angle, *q*, swept out by the two different line segments. (5) Calculate $\cos^2\theta$ and bin this value based on the distance between bead *i* and the condensate centre-of-mass. (6) Repeat steps 1-5 for all chains in the system. (7) For each bin (as described in step 5), we report the average of the calculated $\cos^2\theta$ values. If a given datapoint had an error greater than 0.1, then that datapoint was omitted.

**Statistics and reproducibility**

Fig. 1 and Ext. data figures 1, 2, 4 and 8 show representative images. Other condensates or places in the sample (>50) were observed to show similar architectures. Fig. 2c, d contain information from over 10000 samples (~100pL each). The fitted phase boundaries match with phase boundaries observed for larger samples (10 – 100 µL, Fig. 1). Fig. 3, 4 and Ext. data figures 5-7 show the mean across three replicates and error bars indicating the standard errors. Fig. 5b, d, e shows data collected by imaging a poly-rA condensate over 2.5 s. Fig. 5g shows the results from 6 condensates for 3 different interfaces and 2 interiors, analysed similar to shown in 5b-e. Response table 2 shows how many displacements were analysed for each of these events (Ext. Data Fig. 8). Single molecules blinking at the interface was observed in >50 condensates in all samples.

| Interface | Condensate | # of displacements studied |
|---|---|---|
| rA-dil | 1 | 14446 |
| | 2 | 7182 |
| | 3 | 8881 |
| | 4 | 5601 |
| | 5 | 13628 |
| | 6 | 11091 |
| rA-rC-dil | 1 | 11925 |
| | 2 | 18809 |
| | 3 | 12981 |
| | 4 | 5044 |
| | 5 | 8226 |
| | 6 | 15859 |
| rA-rC | 1 | 4340 |
| | 2 | 7436 |
| | 3 | 6866 |
| | 4 | 3847 |
| | 5 | 6448 |
| | 6 | 2741 |

**Table 1. Number of displacements studied for each datapoint in Fig. 5g.**
